# Overcoming extensive redundancy in the arabidopsis *TREHALOSE-6-PHOSPHATE PHOSPHATASE* gene family reveals connections to development and iron homeostasis

**DOI:** 10.1101/2025.07.09.663849

**Authors:** Tara Skopelitis, Kyle W Swentowsky, Alexander Goldshmidt, Regina Feil, John E. Lunn, David Jackson

**Affiliations:** Cold Spring Harbor Laboratory, Cold Spring Harbor, NY, USA; Institute of Plant Sciences, ARO, Volcani Institute, Rishon LeZion, Israel; Max Planck Institute of Molecular Plant Physiology, 14476 Potsdam-Golm, Germany

## Abstract

Arabidopsis encodes ten *TREHALOSE-6-PHOSPHATE PHOSPHATASE* (*TPP*) genes, homologous to maize *RAMOSA3* (I), which controls shoot branching. To explore the roles of the arabidopsis *TPPs*, we analyzed their expression in shoot apices and found distinct spatial patterns, including *TPPI* and *TPPJ* expressed in shoot meristem boundaries, reminiscent of *RA3* expression. Single and double *TPP* mutants lacked dramatic phenotypes, however a CRISPR-Cas9 knockout of all ten *TPP* genes resulted in increased branching, mirroring *ra3* mutants in maize, as well as reduced size and earlier flowering. Expression of GFP-tagged *TPPI* under its native promoter partially complemented these defects, with protein localization in meristems, vascular tissues and in nuclei. Metabolite profiling revealed higher trehalose 6-phosphate (Tre6P), lower trehalose, and altered sugar and iron-associated metabolites. The mutants also developed chlorosis and grew poorly on low-nutrient media, linked to low iron levels, and reversible with iron supplementation. Consistent with these findings, developmental and iron-responsive genes were up-regulated in the mutants, while photosynthesis-related genes were repressed. Our findings suggest that *TPP* genes redundantly regulate shoot architecture, sugar metabolism, iron homeostasis and photosynthesis in arabidopsis, and support a role for TPP-mediated Tre6P signaling in coordinating developmental and physiological pathways.

## Introduction

Trehalose metabolism plays a critical role in plant development and stress responses, with the intermediate, trehalose 6-phosphate (Tre6P) acting as a key signaling molecule that integrates sugar status with growth regulation (Paul, Gonzalez-Uriarte et al. 2018, Fichtner and Lunn 2021). Trehalose is synthesized by a two-step pathway; trehalose-6-phosphate synthases (TPSs) convert glucose 6-phosphate and UDP-glucose into Tre6P, followed by trehalose-6-phosphate phosphatases (TPPs), which dephosphorylate Tre6P to form trehalose (Cabib and Leloir 1958, Avonce, Mendoza-Vargas et al. 2006, Vandesteene, López-Galvis et al. 2012). Trehalose is present at low levels in most plants, and the intermediate Tre6P functions as a central regulator linking sucrose availability to meristem activity, developmental transitions, and metabolic reprogramming (Paul, Gonzalez-Uriarte et al. 2018, Fichtner and Lunn 2021).

Many plants encode families of *TPS* and *TPP* genes, including 10 arabidopsis *TPP* genes that have both functional redundancy and specialized expression patterns (Leyman, Van Dijck et al. 2001, Vandesteene, López-Galvis et al. 2012). In maize, the *TPP* genes *RAMOSA3* (*RA3*) and *TPP4* control inflorescence shoot branching architecture by controlling meristem determinacy, a function that is likely conserved between species (Satoh-Nagasawa, Nagasawa et al. 2006, Claeys, Vi et al. 2019, Klein, Gallagher et al. 2022). Recent studies have further expanded our understanding of *TPP* gene functions in arabidopsis. For instance, *TPPE* is induced by abscisic acid (ABA) and participates in ABA-mediated root growth and stomatal movement by promoting reactive oxygen species accumulation (Wang, Chen et al. 2020). Tre6P signaling has also been implicated in lateral root formation, acting through Sucrose non-fermenting 1-related protein kinase 1 (SnRK1) and Target of Rapamycin (TOR) signaling in response to auxin (Morales-Herrera, Jourquin et al. 2023) and in shoot meristem function (Lopes, Formosa-Jordan et al. 2024). Mis-expression of heterologous TPPs also impacts shoot branching and flowering (Schluepmann, Pellny et al. 2003), and Tre6P appears to act locally in axillary buds through interaction with strigolactone signaling, as well as systemically through FLOWERING LOCUS T (FT) (Wahl, Ponnu et al. 2013, Fichtner, Olas et al. 2020, Fichtner, Humphreys et al. 2024). One potential mechanism by which Tre6P regulates growth is by binding and inhibiting KIN10, a catalytic subunit of the SnRK1 metabolic sensor (Zhai, Keereetaweep et al. 2018, Blanford, Zhai et al. 2024), or via the interaction of class II TPS proteins with SnRK1 (Van Leene, Eeckhout et al. 2022). These findings suggest that trehalose-related genes integrate metabolism, development, and environmental signaling, and Tre6P has been suggested as a regulatory nexus for carbon signaling in plants (Zhang, Primavesi et al. 2009, Figueroa and Lunn 2016, Baena-Gonzalez and Lunn 2020) A comprehensive understanding of the biochemical roles, regulatory networks and spatial expression patterns of the *TPP* gene family could therefore improve crops through targeted metabolic and developmental engineering. Here we knockout all ten arabidopsis *TPP* genes using CRISPR-Cas9, and dissect their roles in development and physiology, revealing connections to iron homeostasis, sugar metabolism and development.

## Results

The arabidopsis genome encodes a family of ten *TPP* genes. To identify potential members involved in inflorescence development, analogous to *RA3* function in maize (Satoh-Nagasawa, Nagasawa et al. 2006, Claeys, Vi et al. 2019), we first characterized their expression patterns in shoot apices using a combination of *in situ* hybridization and β-glucuronidase (GUS) staining. The ten *TPP* genes had diverse expression patterns (Fig. 1), with several having diffuse or undetectable staining, likely reflecting low or constitutive expression. However, some had more localized expression, including *TPPG* expressed in the center of the inflorescence meristem and in the expanding stem below, as evidenced by a GUS gene trap line, and TPPF, expressed throughout the inflorescence and floral meristems. Two of the genes, *TPPI* and *TPPJ*, were expressed in meristem boundaries (arrowed, Fig. 1I, J). These patterns were reminiscent of the *RA3* expression pattern in maize, in arcs of cells at the base of spikelet pair and spikelet meristems (Satoh-Nagasawa, Nagasawa et al. 2006, Claeys, Vi et al. 2019).

**Fig. 1.**
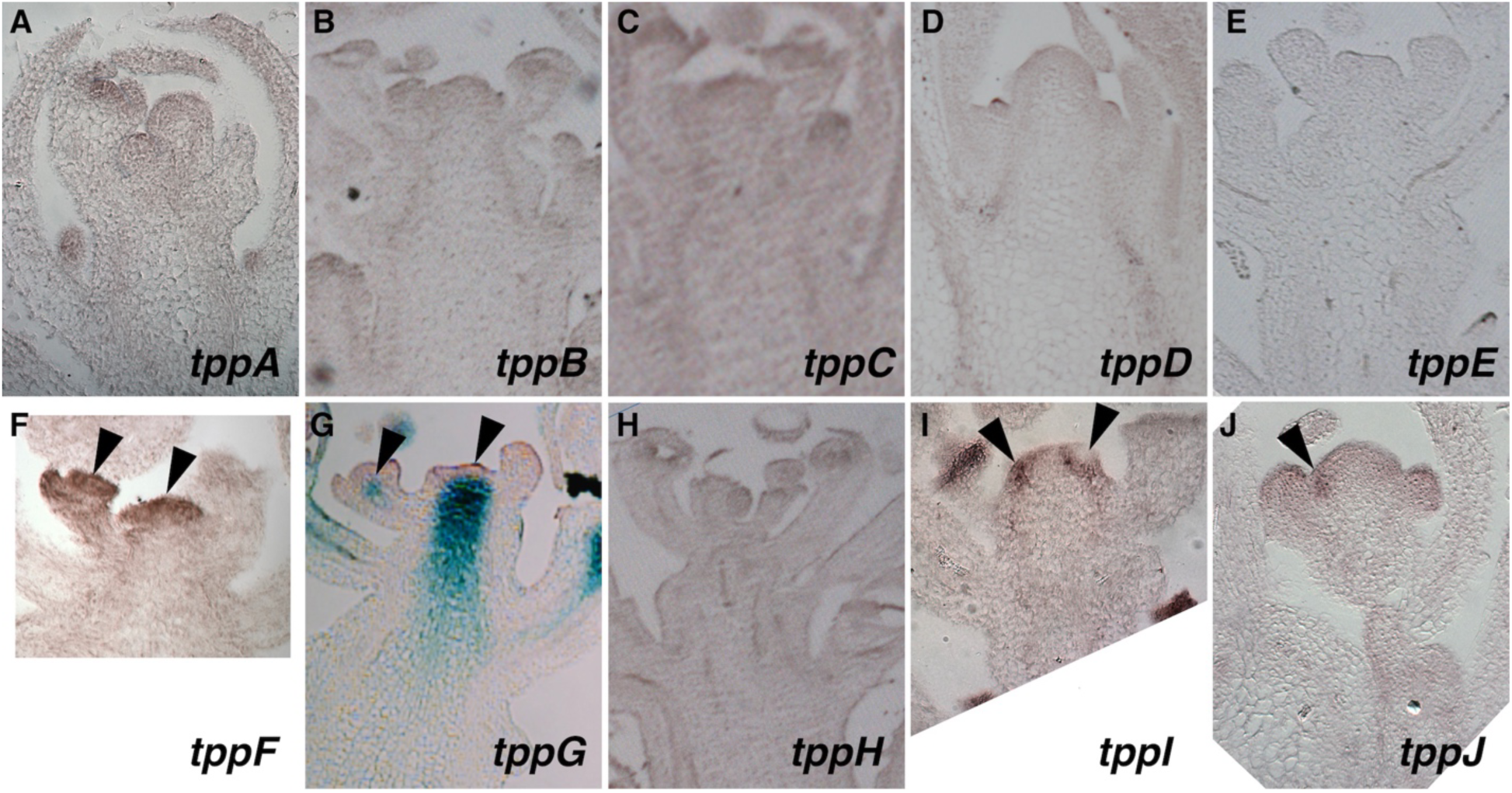
Arabidopsis *TPP* genes have diverse expression patterns. Gene expression was detected by *in situ* hybridization, except for TPPG, where a gene trap line was used. Specific staining is indicated by black arrows. In panel G, TPPG expression (blue) is evident in the central region of inflorescence and floral meristems, arrowed. In panel F, *TPPF* expression is throughout inflorescence and floral meristems. In panels I and J, TPPI and TPPJ expression is observed in boundary domains between the inflorescence and floral meristems, arrowed.

Based on their conserved expression pattern, we knocked out *TPPI* and *TPPJ* using CRISPR-Cas9 genome editing, however the double mutant plants lacked striking developmental phenotypes. To overcome possible redundancy from additional family members, we therefore knocked out all 10 *TPP* genes, using a multiplex CRISPR strategy. We designed two constructs, each targeting four of the *TPP* genes with two guide RNAs each, and transformed one construct into wild type (Columbia-0, col) and the other into the *tppi; j* double mutant. Plants were genotyped and ones carrying mutations in the four newly targeted genes were crossed together. Next, the selfed progeny were genotyped to identify plants with homozygous or biallelic mutations in all 10 *TPP* genes. The 10x *tpp* knockout plants were grown alongside wild-type plants for phenotyping.

The 10x *tpp* knockout plants were significantly smaller, with a mature rosette diameter around half of the wild type plants (Fig. 2), and a mean plant height around 2/3 of the wild type plants under long day conditions. Similar results were found for plants grown in short days (Fig. 2). The knockout plants also flowered earlier, after ∼20 days in long days, compared to ∼26 days for the wild type plants, and after ∼55 days in short days, compared to ∼75 days. Similar to maize *ra3* mutants, the 10x *tpp* ko mutants were also significantly more branched, with ∼11 primary rosette branches per plant, compared to ∼6 for the wild type plants grown in long days, and enhanced primary rosette branching was also observed in short days (Fig. 2).

**Fig. 2.**
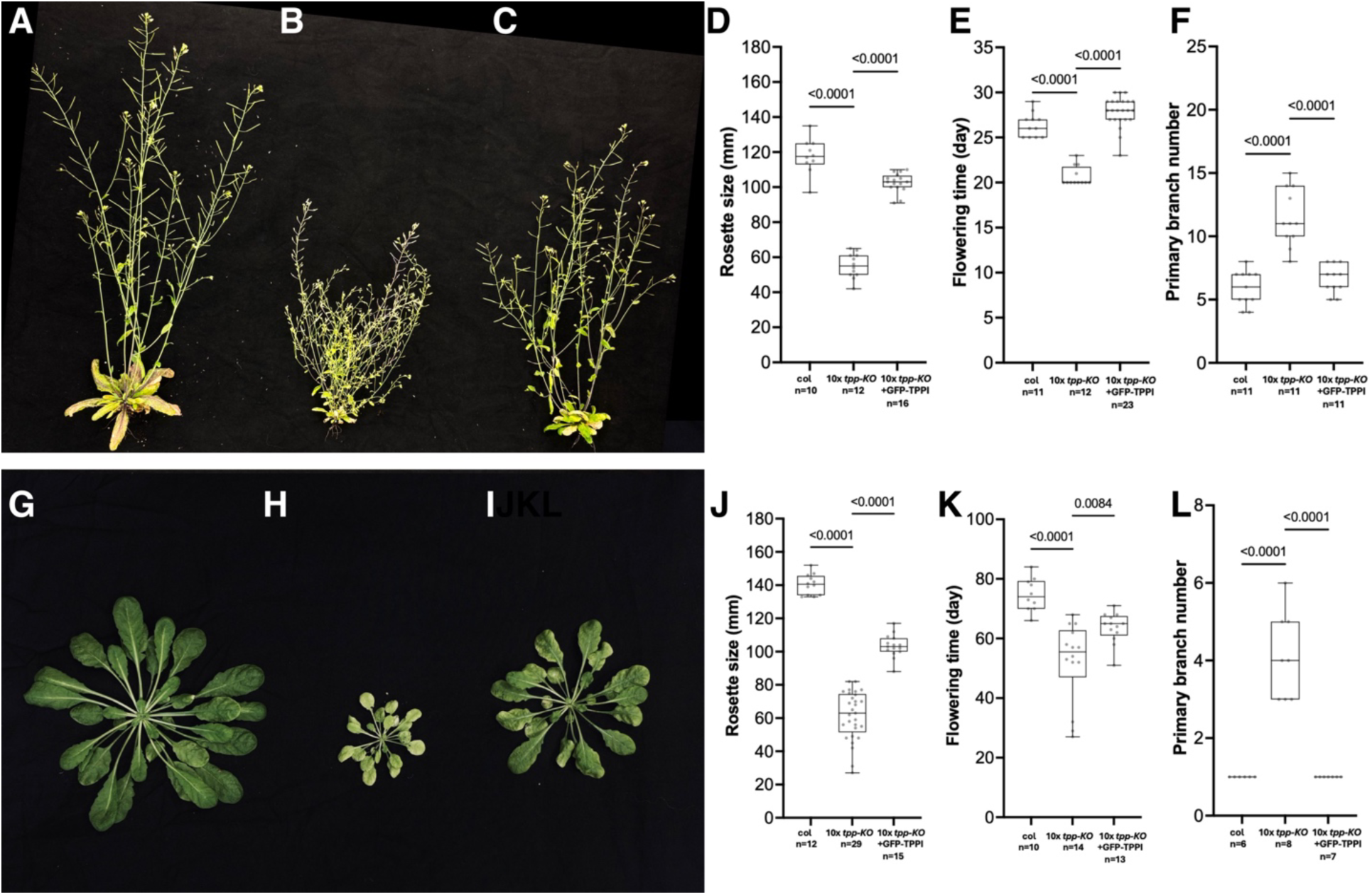
10x *tpp* knockout plants are smaller with enhanced branching. Wild type (Col) plants (A, G), 10x *tpp* knockout plants (B, H) and 10x *tpp* knockout plants complemented with native promoter driven *GFP-TPPI* (C, I) were grown under long days (A-C) or short days (G-I). The graphs show corresponding quantification of rosette size, flowering time and branch number for wt (col), 10x *tpp-*knockout *(KO)* and 10x *tpp-KO* complemented with pTPPI-GFP-TPPI. The box plots show individual measurements as dots, each box represents the middle 50% of the data, with the line inside marking the median and the edges of the box show the lower and upper quartiles, while whiskers extend to the minimum and maximum values. P-values are as marked.

We next sought to understand the spectrum of redundancy in *TPP* genes, by asking if the 10x *tpp* ko phenotypes could be complemented by expression of a single *TPP* gene. We designed a construct based on the meristem-expressed *TPPI* genomic sequence, including 2.1kb promoter, introns and 3’ sequences, to express a chimeric GFP-TPPI fusion protein, and indeed could partially complement the knockout phenotypes. We generated 14 independent transgenic lines, and selected one line, #11, with a rosette size close to wild type, for further characterization. Native expression of GFP-TPPI partially complemented all aspects of the 10x *tpp* knockout phenotypes, including rosette size, plant height and branching, in both short and long day conditions (Fig. 2). Confocal microscopy revealed that GFP-TPPI was expressed on the flanks of floral meristems (Fig. 3B), similar to our *in situ* hybridization results for the endogenous *TPPI* transcript (Fig. 1I). Consistent with this meristem enriched expression, the 10x *tpp* knockout mutants had smaller shoot meristems (Fig. 3G). We also observed GFP-TPPI expression in vascular strands, likely in the phloem, as expression was in cells adjacent to the distinctive xylem vessels (Fig. 3C). Natively expressed GFP-TPPI was also observed in roots, predominantly in the outer layer of the stele (Fig. 3E), and in initiating lateral root primordia (Fig. 3F). In all tissues, GFP-TPPI appeared to accumulate in nuclei, and localized to sub-nuclear puncta (arrowed, Fig. 3D), reminiscent of RA3 localization in maize (Demesa-Arevalo, Abraham-Juarez et al. 2021).

**Fig. 3.**
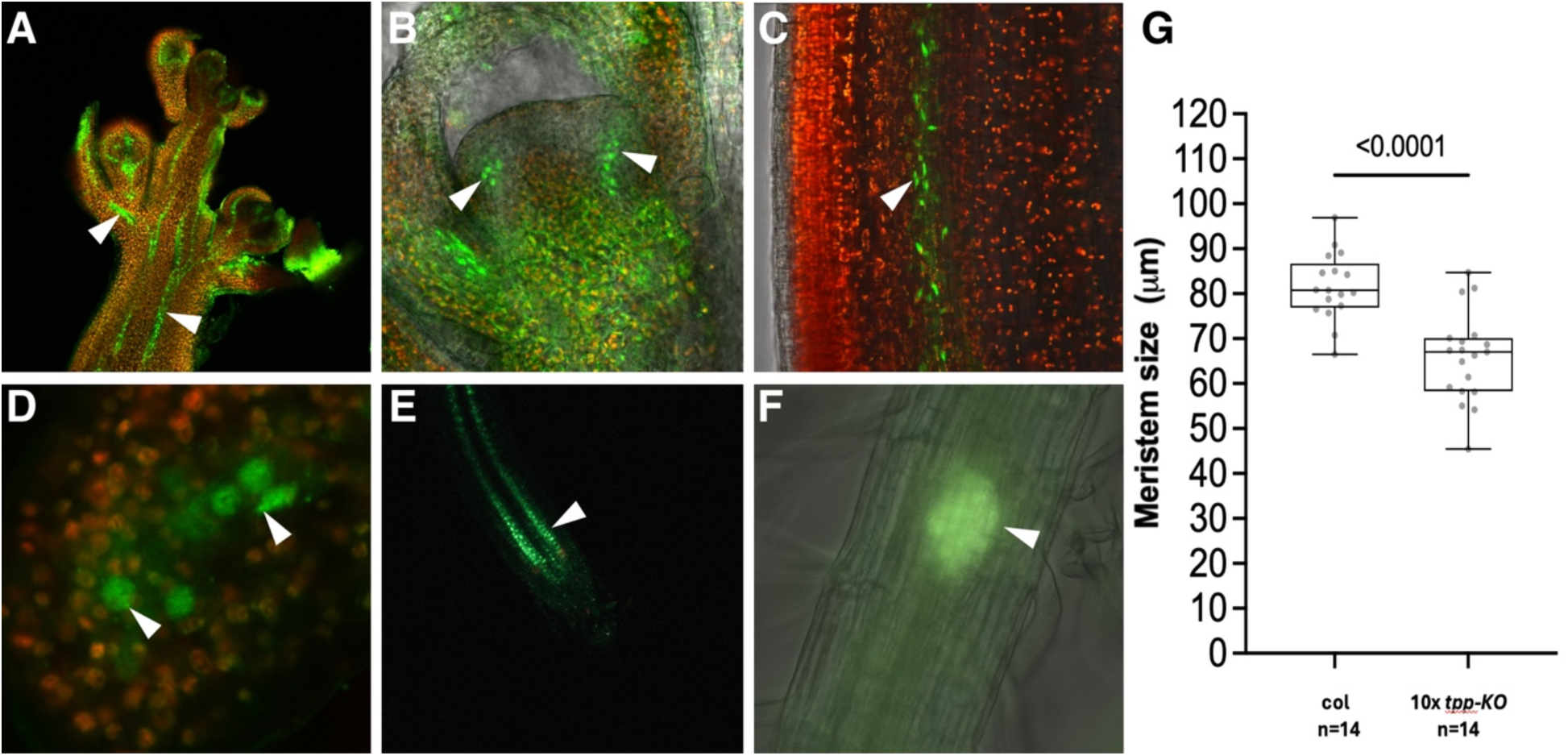
A native promoter driven GFP -TPPI fusion is expressed in meristem boundaries and vascular tissues. (A) inflorescence apex, vascular strands arrowed, (B) Close up of floral meristem, with expression on flanks of meristem arrowed, (C) Inflorescence stem close up, vascular expression arrowed, (D) Close up of leaf primordium cells, GFP-TPPI localizes to nuclei, and nuclear puncta, arrowed, (E) expression in root tip, localized to outer layer of stele (F) GFP- TPPI expression in initiating lateral root primordium, arrowed. In all images, GFP-TPPI expression is green or yellow, chlorophyll autofluorescence is red. Consistent with meristem expression, the 10x *tpp* knockout plants have smaller shoot meristems (G). The box plot shows individual measurements as dots, each box represents the middle 50% of the data, with the line inside marking the median and the edges of the box show the lower and upper quartiles, while whiskers extend to the minimum and maximum values.

To understand the physiological effects of mutating all arabidopsis *TPP* genes, we next measured sugar-related metabolites. We measured two sets of pooled rosette samples, one collected just before the end of the night period, and one collected just before the end of the day, with six biological replicates. Tre6P levels were higher at the end of the day than at the end of the night, as expected, reflecting higher sucrose levels at the end of the day (Fig. 4). Also as expected from loss of all *TPP* genes, Tre6P levels were significantly higher in the 10x *tpp* knockout mutants (Fig. 4), and trehalose levels were significantly lower. Sucrose levels trended lower in the end of night samples, though the differences were not significant, and sucrose:Tre6P ratios trended lower, though only significant in end of night samples. Raffinose, and precursors myo-inositol and galactinol, also trended higher in the knockout mutants, suggesting a stress response (Yan, Qing et al. 2022). We also found a >100-fold difference in ADP glucose (ADPG) levels between day and night samples. ADPG turns over rapidly, and is extremely sensitive to shading, suggesting our sampling accurately preserved the *in vivo* metabolite levels (Fünfgeld, Wang et al. 2022).

**Fig. 4.**
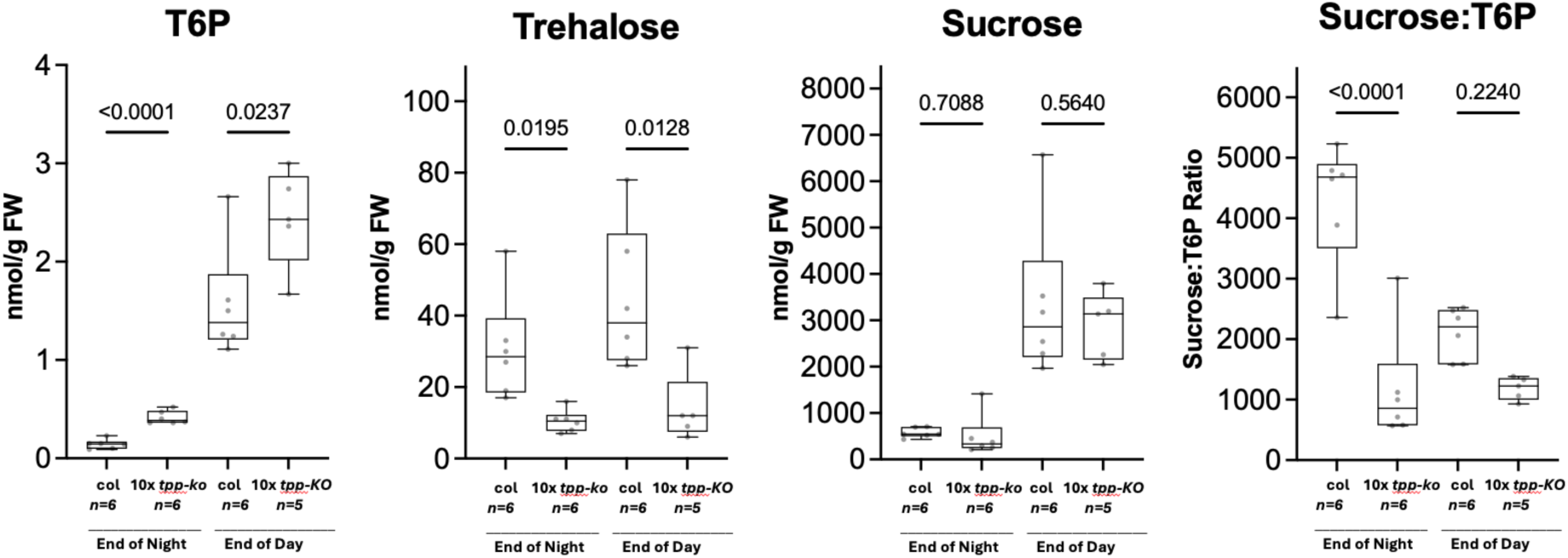
10x *tpp* knockout plants have higher T6P levels and lower trehalose. The plots show measurements of the indicated sugars in wt (columbia, col) and 10x *tpp* knockout mutants just before end of night and end of day, as marked. The box plots show individual measurements as dots, each box represents the middle 50% of the data, with the line inside marking the median and the edges of the box show the lower and upper quartiles, while whiskers extend to the minimum and maximum values. P-values are as marked.

In addition to the morphological and sugar metabolite phenotypes, the 10x *tpp* knockout plants were significantly paler than normal, with a yellowish appearance, especially when grown on media low in nutrients. For example, the mutants were very pale when grown on ¼ strength Murashige-Skoog (MS) media plates (Fig. 5A, B). Again, this phenotype was largely complemented by expression of natively expressed GFP-TPPI (Fig. 5C). The 10x *tpp* knockout plants continued to grow slowly when transferred to soil, and in the absence of supplemental fertilizer were extremely pale and produced flowers, but failed to make seed. However, the plants became green within a few days after adding fertilizer, and could set seed (Fig. 5D-F). To further understand this phenotype, we measured different metal elements by acid extraction of dried, ground and ashed material (J.R. Peters). The levels of several metal micronutrients were altered in the 10x *tpp* knockout mutants, with significantly lower iron levels (Fig. 5G). We therefore also attempted to rescue the pale phenotype using a solution of iron and other metal micronutrients, (“liquid iron”; Garden Rich), and this was also sufficient to rescue the pale phenotype and restore green color. The 10x *tpp* ko mutants also had a severely reduced root system, and defects in root hair development, including higher density and abnormal branching of root hairs.

**Fig. 5.**
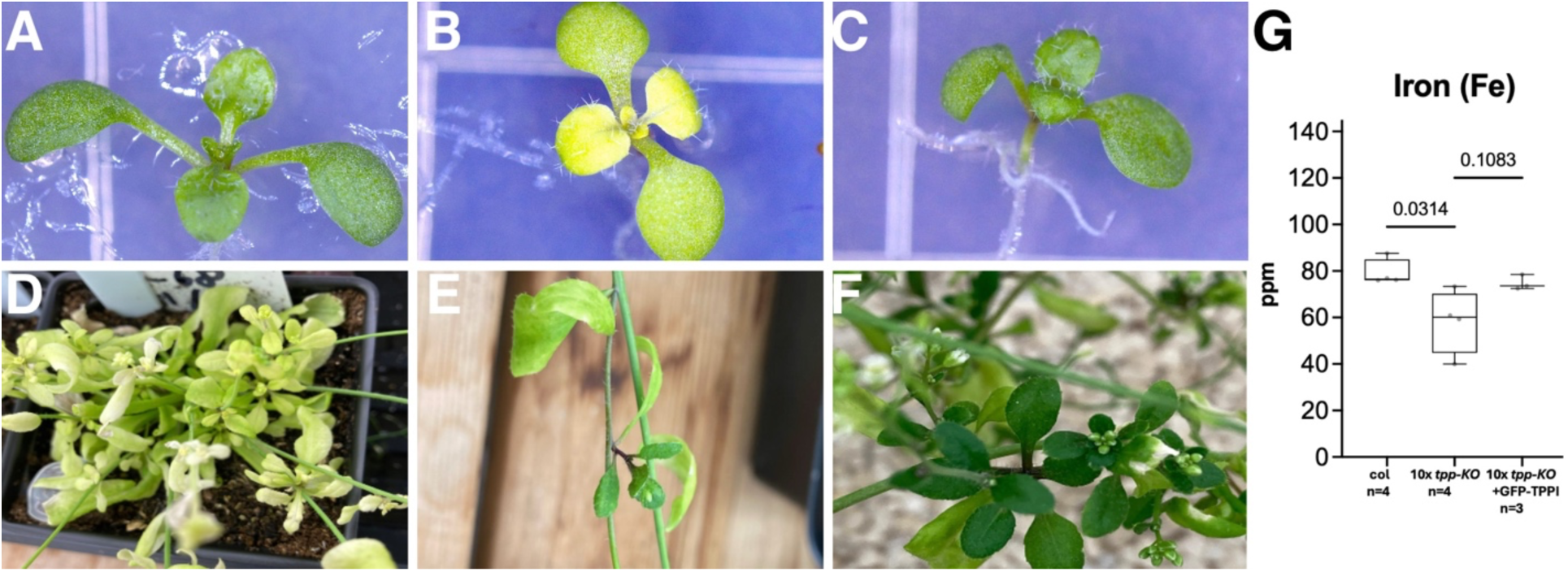
10x *tpp* knockout plants were pale, and rescued by iron. Seedlings of Columbia wild type (A), 10x *tpp* knockout plants (B) and 10x *tpp* knockout plants complemented with native promoter driven *GFP-TPPI* (C) grown on ¼ MS- media, the 10x tpp knockout mutants are yellowish. Later in development, the 10x *tpp* knockout mutants continue to be pale in the absence of fertilizer (D) but new growth is green a few days after fertilizer application (E, F). (G) Iron levels were significantly lower in the 10x tpp knockout mutants compared to columbia (col) wild type and partially restored in 10x *tpp* knockout plants complemented with native promoter driven *GFP-TPPI.* The box plots show individual measurements as dots, each box represents the middle 50% of the data, with the line inside marking the median and the edges of the box show the lower and upper quartiles, while whiskers extend to the minimum and maximum values. P-values are as marked.

To understand the impact of the 10x *tpp* knockouts on developmental gene expression, we next dissected inflorescence apices for mRNA sequencing. Approximately 1,600 genes were significantly upregulated in the 10x *tpp* knockout mutants, and ∼660 genes were downregulated (log2 fold change > 1 or < -1). Three of the *TPP* genes, *TPPF*, *TPPG* and *TPPJ* were upregulated in the 10x *tpp* knockout mutants, with *TPPJ* upregulated ∼ 3.3 fold, suggesting it actively compensates for loss of *TPP* expression, similar to the interaction between *RA3* and *TPP4* paralogs in maize (Satoh-Nagasawa, Nagasawa et al. 2006, Claeys, Vi et al. 2019). Gene ontology (GO) analysis found significant enrichment in upregulated genes related to development, including “primary shoot apical meristem specification” (q-value=1.20 E-3), and “developmental growth involved in morphogenesis” (q-value =2.69 E-3) (Fig. 6A). Upregulated developmental genes included meristem boundary gene *CUP SHAPED COTYLEDONS2* (*CUC2*), leucine rich repeat receptor genes *RECEPTOR-LIKE PROTEIN KINASE 2* (*RPK2*), *BARELY ANY MERISTEM1* (*BAM1*), auxin transporters *PINFORMED2* (*PIN2*) and *LIKE AUXIN RESISTANT1* (*LAX1*), chromatin regulator *TOPLESS* (*TPL*) and sugar transporters *SUGARS WILL EVENTUALLY BE EXPORTED TRANSPORTER* (*SWEET*) *3, 13, 14* and *15*. Downregulated development associated genes included *SMALL AUXIN UPREGULATED RNA* (*SAUR*) *4*, *14*, *16*, *49* and *50*. Canonical shoot meristem genes, such as *WUSCHEL* (*WUS*) and *CLAVATA* (*CLV*) *1*, *2* and *3* were not differentially expressed, nor was the branching regulator *BRANCHED1* (*BRC1*). However, multiple hormone related pathways were mis-regulated. Several upregulated GO categories were related to iron, including “response to iron ion” (q-value= 1.13 E-2) and “iron ion transport” (q-value= 4.03 E-4) (Fig. 6A). A GO category “phloem transport” was also enriched (q-value= 4.10 E-2), consistent with vascular-enriched expression of *GFP-TPPI.* “Positive regulation of transcription elongation by RNA polymerase II” was also enriched, consistent with the localization of RA3 and GFP-TPPI to nuclear puncta (Demesa-Arevalo, Abraham-Juarez et al. 2021).

**Fig. 6.**
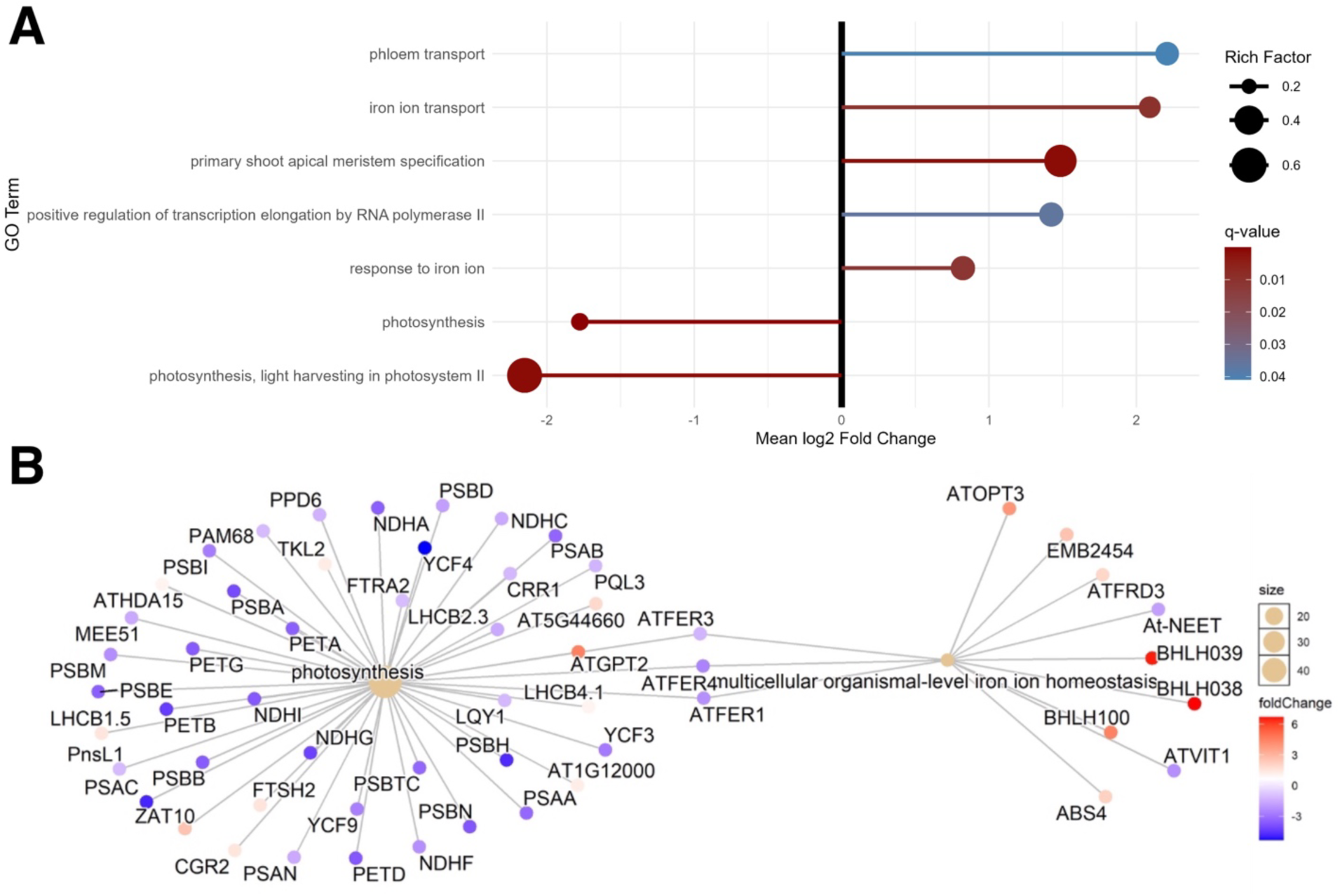
mRNAseq analysis of 10x *tpp* knockout plants identifies GO terms related to development, iron and photosynthesis. (A) Plot of selected GO terms, line lengths represent mean fold change, ball size and color reflect proportion of DEGs compared to all genes in a given GO category, and adjusted p-value, respectively.

Many of the downregulated genes in 10x *tpp* knockout plants were related to photosynthesis, including GO categories “photosynthesis” (q-value= 2.75 E-8) and “photosynthesis, light harvesting in photosystem II” (q-value= 1.21 E-3) (Fig. 6A). The downregulation of photosynthesis related GO terms and upregulation of iron related terms is consistent with our observation that the 10x *tpp* knockout plants were pale and rescued by iron treatment. A network analysis indicated that the DEGs related to iron and photosynthesis may be connected through arabidopsis *FERREDOXIN* (*AtFER*) genes (Fig. 6B). Furthermore, KEGG integration of our sugar metabolite and mRNAseq analysis revealed potential regulatory points in starch and sucrose metabolism affected in the 10x *tpp* knockout plants.

In summary, we present an analysis of arabidopsis plants lacking all *TPP* genes. The mutants were more branched, reminiscent of maize plants with knockouts of one or two TPP family members, and were smaller and flowered earlier, reminiscent of transgenic arabidopsis plants with elevated Tre6P (Fichtner et al., 2021). The knockout plants also had defects in photosynthesis and iron metabolism, supporting the central importance of *TPP* genes and Tre6P signaling in plant development and metabolic regulation.

## Discussion

*TPP* genes encode enzymes that dephosphorylate the signaling molecule Tre6P to trehalose, and are important regulators of plant development and physiology. Many studies have examined the role of Tre6P signaling by mis-expression of TPP or TPS enzymes from bacterial or other sources (reviewed in (Fichtner and Lunn 2021, Miret, Griffiths et al. 2024), and indicate roles for Tre6P in control of root branching (Morales-Herrera, Jourquin et al. 2023) and shoot branching (Fichtner, Barbier et al. 2017, Fichtner, Barbier et al. 2021), among other developmental and physiological phenomena (Wahl, Ponnu et al. 2013, Lopes, Formosa-Jordan et al. 2024). However, few studies investigated the endogenous roles of *TPP* genes using loss of function mutations, presumably because of complications due to extreme redundancy. Expression of arabidopsis *TPP* family members differs at cell-, tissue- and developmental stage-levels, and all 10 genes encode active TPP enzymes (Vandesteene, López-Galvis et al. 2012), suggesting they exert spatial control over Tre6P levels and signaling (reviewed in (Kerbler, Armijos-Jaramillo et al. 2023).

A small number of *TPP* genes have been characterized by single or double mutant analyses. For example, *tppG* loss-of-function mutants are less sensitive to abscisic acid (ABA) in root inhibition and stomatal closure (Vandesteene et al., 2012) as are *tppE* mutants (Wang, Chen et al. 2020), while *tppF* loss-of-function mutants are drought-sensitive (Lin, Yang et al. 2019). In contrast, *tppI* mutants are sensitive to chilling stress (Lin, Wang et al. 2023), while *tppD* mutants are hypersensitive to salinity stress (Krasensky, Broyart et al. 2014). Similar results were found in rice, where *tpp1* mutants have higher ABA levels and slow germination (Wang, Li et al. 2021), *tpp3* mutants are less salt tolerant (Ye, Wang et al. 2023) and *OsTPP7* promotes anaerobic germination (Kretzschmar, Pelayo et al. 2015). Of note, our 10x *tpp* knockout plants accumulated higher levels of sodium ions, perhaps related to a function in salinity stress (Zhou, Shi et al. 2024).

Limited developmental phenotypes of *tpp* mutants have also been described. For example, *tppB* mutants have an increase in cell number and larger leaves, and *TPPB* acts downstream of auxin to control lateral root formation (Morales-Herrera, Jourquin et al. 2023). *tppI* mutants have defects in primary and lateral root elongation and auxin transport (Lin, Wang et al. 2020), and *tppJ* mutants flower early through mis-regulation of *SQUAMOSA-PROMOTER BINDING PROTEIN-LIKE* (*SPL*) genes (Musialak-Lange, Fiddeke et al. 2021). One indicator of potential redundancy in the *TPP* gene family comes from studies of root hair formation, since *tppA; tppG* double mutants make additional root hairs, while the single mutants appear normal (Van Houtte, Lopez-Galvis et al. 2013). However, up to now, a role of plant *TPP* genes in shoot branching has only been described in maize, where single mutants of *RA3* have ectopic ear branches and additional tassel branches, and double mutants with its’ paralog *TPP4* are greatly enhanced (Satoh-Nagasawa, Nagasawa et al. 2006, Claeys, Vi et al. 2019). Here we found that shoot branching is enhanced in arabidopsis plants with knockouts of all 10 *TPP* genes, similar to the maize single or double mutants. Differential redundancy between species is a common phenomenon, highlighting the benefit of studying different model systems (Láruson, Yeaman et al. 2020), and our results suggest that control of branching by *TPP* genes is universal in angiosperms.

The 10x *tpp* knockout plants were also stunted, and flowered earlier, consistent with studies of conditional TPS expression (Schluepmann, Pellny et al. 2003, Wahl, Ponnu et al. 2013, Fichtner, Olas et al. 2020). Metabolite profiling found, however, that the 10x *tpp* knockout plants had only a relatively modest increase in Tre6P levels, and trehalose levels were not reduced to zero. These observations suggest that plants have cryptic TPP activity from non-specific phosphatases or low-level, promiscuous activity of more specific phosphatases, such as sucrose-6^F^-phosphate phosphatase. The latter catalyzes the dephosphorylation of sucrose 6^F^-phosphate, the intermediate of sucrose biosynthesis analogous to Tre6P. This enzyme can bind trehalose in the active site (Fieulaine, Lunn et al. 2007), so might also be able to dephosphorylate Tre6P, albeit at low efficiency. Another possibility is that the class II TPSs have cryptic TPP activity. These proteins lack TPS activity, but have TPP-like domains with almost perfectly conserved active sites that have been retained throughout the evolution of the green plant lineage from chlorophyte algae to angiosperms (Lunn 2007, Vandesteene, Ramon et al. 2010). Despite this, the plant class II TPSs are unable to complement the yeast *tps2Δ* mutant that lacks TPP activity (Ramon, De Smet et al. 2009), and recombinant class II TPS proteins lack TPP activity *in vitro* (Harthill, Meek et al. 2006).

However, we cannot exclude the possibility that the recombinant proteins lack activity due to misfolding, or they might lack essential post-translational modifications when expressed in yeast or *Escherichia coli*. Therefore, it is possible that the class II TPSs have TPP activity *in planta*, and this might only be detectable when the canonical TPPs are missing, as in our 10x *tpp* mutant.

An unexpected finding from the 10x *tpp* knockout plants was their pale phenotype, iron deficiency and significant mis-expression of iron and photosynthesis related genes. Our results reveal a previously unknown connection between Tre6P and iron signaling. Iron deficiency can result from multiple causes, including defective uptake, signaling or response (Vélez-Bermúdez and Schmidt 2021, Liang 2022). The 10x *tpp* knockout plants had striking defects in root growth and root hair patterning, suggesting uptake as a possible cause of iron deficiency. Another possibility is a defect in exudation of malate and citrate, organic acids used to acidify the rhizosphere to solubilize otherwise immobile cations, especially Fe^2+^and Fe^3+^, since a connection has been made between root exudates and Tre6P (Shane, Feil et al. 2016). Iron related genes are also differentially expressed in *snrk1* mutants and over-expression lines (Peixoto, Moraes et al. 2021), and trehalose related gene expression and Tre6P levels correlate with *SnRK1* activity (Nunes, O’Hara et al. 2013, Avidan, Martins et al. 2024, Blanford, Zhai et al. 2024), supporting the Tre6P-iron connection. Whether specific members of the *TPP* gene family control this effect will require analysis of different knockout combinations.

The 10x *tpp* knockout phenotypes could be largely rescued by native expression of a single *TPP* gene, *TPPI*. Imaging of the complementing GFP-TPPI fusion found expression in shoot meristem boundary domains, in developing vascular tissues and in root stele tissues and lateral root primordia. Expression in vascular tissues is reminiscent of *TPS1,* the single active *TPS* gene in arabidopsis (Fichtner, Olas et al. 2020), and suggests that vascular tissues, meristems and initiating primordia are important sites of Tre6P signaling. We also observed localization of the GFP-TPPI fusion in sub-nuclear puncta, similar to RA3 in maize (Demesa-Arevalo, Abraham-Juarez et al. 2021) supporting the idea that *TPP* genes play a moonlighting function in gene regulation (Claeys, Vi et al. 2019). Our results also suggest that Tre6P, or a downstream factor, could act as a mobile signal, since the yellow leaf phenotype was largely complemented by GFP- TPPI expression in vascular tissues, whereas yellowing results from changes in photosynthetic mesophyll cells.

In summary, we show the power of CRISPR-Cas9 editing to mutate a complete gene family, and reveal new hypotheses about the role of TPPs and Tre6P in plant development and physiology. Follow up work should tease out the roles of individual TPP members in these traits as well as the significance of nuclear localization of TPP enzymes.

## Materials and methods

### Plant growth and manipulation

Seeds were sown on either full, 1/2, or ¼ Murashige and Skoog (MS) medium after vapor-phase sterilization (Lindsey, Rivero et al. 2017). Seeds were stratified at 4°C in the dark for 2–4 days to synchronize germination, and transferred to growth chambers. Plants were transferred to soil (Premier Tech Pro - Mix FPX Biofungicide), and grown under either short-day- 8-hour light/16-hour dark cycle or long-day- 16-hour light/8-hour dark cycle, and watered with constant rate 50ppm nitrogen Jack’s Professional 20-20-20 general-purpose fertilizer with micronutrients, or no fertilizer. Light was provided at an intensity of approximately 120–150 μmol m⁻² s⁻¹, and temperature was maintained at 22°C during the light period and 18–20°C during the dark period, with relative humidity between 50–70%. For iron treatments, plants were grown in soil in a growth chamber and treated with Garden Rich Liquid Iron + Micronutrients containing 5% ferrous sulfate, a non-chelated Fe^2+^, applied as a soil drench at recommended concentrations. Pale plants were treated twice, one week apart.

CRISPR-Cas9 gene editing was performed as described (Nimchuk 2017), using primers listed in table S3.

A GFP-TPPI fusion construct was generated using 2kb *TPPI* promoter upstream of the translational start (ATG) and *TPPI* coding sequence (∼3.5kb) and *GFP* sequences cloned into pBA002a vector (Sanchez, Ullman et al. 2002) using primers listed in table S3. Plasmids were transformed into Arabidopsis using floral dip protocol as described (Clough and Bent 1998) using agrobacterium strain GV3101.

### Meristem Measurements

Arabidopsis shoot apices were dissected from 17 day old plants grown in short-day conditions, and fixed overnight at 4 °C in FAA solution (3.7% formalin, 5% acetic acid, and 50% ethanol in water), dehydrated through an ethanol series, then cleared through increasing concentrations of Histo-Clear II (National Diagnostics for imaging). After mounting on slides, the cleared were photographed using 20x magnification on a Nikon Model Eclipse Ni-E light microscope and measured using ImageJ software.

#### *in situ* hybridization and gene trap line staining

*in situ* hybridization was performed using probes obtained by PCR from arabidopsis cDNA with primers flanking cDNA sequences unique for each one of the arabidopsis *TPP* genes, except *TPPG*, with T7 RNA polymerase promoter incorporated at the reverse primer. The resulting PCR products were used for *in vitro* transcription with DIG-labeled UTP for probe synthesis and *in situ* hybridization, as described by (Jackson, Veit et al. 1994).

GUS staining of *TPPG* (*At4g22590*) gene trap line GT24630 containing a GUS insertion in the same orientation was performed as described (McConnell and Barton 1998) with 90% acetone fixation and the addition of 20% methanol into the staining solution to reduce non-specific background.

### Confocal imaging

Arabidopsis shoot apices were embedded in 6% low-melting-point agarose and sectioned using a vibratome (Leica VT1000S) at 100 µm thickness. Sections were mounted in water on glass slides, and confocal imaging performed using a Zeiss LSM 910 laser scanning confocal microscope. Roots were mounted directly on slides. Fluorescent signals were detected using appropriate laser lines and emission filters depending on the fluorophore(s) used (e.g., 488 nm for GFP, 561 nm for chlorophyll). Image acquisition settings (e.g., laser power, gain, and pinhole size) were optimized to minimize photobleaching and ensure consistent signal detection across samples. Image processing was performed using Zeiss ZEN software and Fiji (ImageJ).

### RNAseq and analysis

Inflorescence apices were dissected to enrich for meristematic and young floral tissues, excluding mature flowers. Tissue was collected in biological triplicates, flash-frozen in liquid nitrogen, and total RNA was extracted using the RNeasy Plant Mini Kit (Qiagen) with on-column DNase I treatment. RNA integrity was confirmed using an Agilent 2100 Bioanalyzer, and RNA quantity was measured using a Qubit fluorometer. RNA-seq libraries were prepared using the Illumina TruSeq Stranded mRNA Library Prep Kit and sequenced on an Illumina NovaSeq 6000 platform to generate ∼30 million 150 bp paired-end reads per sample. Raw reads were quality-checked using FastQC, trimmed with Trimmomatic, and aligned to the *Arabidopsis thaliana* TAIR10 reference genome using STAR. Gene-level counts were quantified with featureCounts, and differential expression analysis was performed using DESeq2 in R. Genes with an adjusted p-value < 0.05 and |log2(fold change)| > 1 were considered significantly differentially expressed.

Gene Ontology (GO) enrichment analysis was conducted using the clusterProfiler R package, with significance determined by adjusted p-values (< 0.05). KEGG pathway enrichment analysis was also performed using clusterProfiler, mapping Arabidopsis genes to KEGG Orthology terms. Visualization of enriched pathways and GO categories was done using ggplot2.

### Metabolite analysis

Tre6P and other metabolites were extracted with chloroform-methanol as described in Lunn et al. (2006). Tre6P, other phosphorylated intermediates and organic acids were measured by high-performance anion-exchange liquid chromatography coupled to tandem mass spectrometry as described by Lunn et al. (2006) with modifications (Figueroa et al., 2016). Sugars were measured by high-performance hydrophilic interaction liquid chromatography coupled to tandem mass spectrometry as described in Reis-Barata et al. (manuscript in preparation).

## Author Contributions

TS performed all experiments and analysis, except as noted below; KWS performed RNAseq and other informatics analyses and made relevant figures, AG performed *in situ* hybridization and GUS staining, RF and JL performed sugar measurements and analysis, DJ conceptualized the research and made figures and wrote the manuscript; JEL co-wrote the manuscript and all authors edited the manuscript.

## Acknowledgements

The study is funded by the NSF award IOS 2131631. R.F. and J.E.L. were financially supported by the Max Planck Society. We thank Thu Tran for assistance with statistical analysis. We thank Ullas Pedmale for the use of a confocal microscope, and the Uplands farm team, including Tim Mulligan, Kyle Schlecht, and Autumn Harrison, for plant care, and Dr. Jae Hyung Lee for construction and transformation of the GFP-TPPI plasmid.

